# Proximity Measures as Graph Convolution Matrices for Link Prediction in Biological Networks

**DOI:** 10.1101/2020.11.14.382655

**Authors:** Mustafa Coşkun, Mehmet Koyutürk

## Abstract

**Motivation:** Link prediction is an important and well-studied problem in computational biology, with a broad range of applications including disease gene prioritization, drug-disease associations, and drug response in cancer. The general principle in link prediction is to use the topological characteristics and the attributes–if available– of the nodes in the network to predict new links that are likely to emerge/disappear. Recently, graph representation learning methods, which aim to learn a low-dimensional representation of topological characteristics and the attributes of the nodes, have drawn increasing attention to solve the link prediction problem via learnt low-dimensional features. Most prominently, Graph Convolution Network (GCN)-based network embedding methods have demonstrated great promise in link prediction due to their ability of capturing non-linear information of the network. To date, GCN-based network embedding algorithms utilize a Laplacian matrix in their convolution layers as the convolution matrix and the effect of the convolution matrix on algorithm performance has not been comprehensively characterized in the context of link prediction in biomedical networks. On the other hand, for a variety of biomedical link prediction tasks, traditional node similarity measures such as Common Neighbor, Ademic-Adar, and other have shown promising results, and hence there is a need to systematically evaluate the node similarity measures as convolution matrices in terms of their usability and potential to further the state-of-the-art.

**Results:** We select 8 representative node similarity measures as convolution matrices within the single-layered GCN graph embedding method and conduct a systematic comparison on 3 important biomedical link prediction tasks: drug-disease association (DDA) prediction, drug–drug interaction (DDI) prediction, protein–protein interaction (PPI) prediction. Our experimental results demonstrate that the node similarity-based convolution matrices significantly improves GCN-based embedding algorithms and deserve more attention in the future biomedical link prediction

**Availability:** Our method is implemented as a python library and is available at githublink

**Supplementary information:** Supplementary data are available at *Bioinformatics* online.

## 1 Introduction

Graphs (networks) are one of the main data science tools to represent biomedical entities (as nodes) and their relations (as edges). Developing computational methods to analyze and understand these networks is one of the major challenges for bioinformatics researchers. One of these research challenges is discovering new interactions (links) in the biomedical networks, i.e., solving link prediction problem in biological networks. A considerable amount of research efforts has been devoted to developing computational methods to predict/identify the missing/spurious links in various biomedical networks, such as drug-disease association (DDA) networks Liang *et al.* (2017), detecting long non-coding RNA (lncRNA) functions on protein interaction graphs Zhang *et al.* (2018), and predicting missing links for drug-response problem on drug and protein networks Stanfield *et al.* (2017).

One ways of tacking the biological link prediction problem is to extract features that represent network topology, graph representation learning techniques are used to embed the nodes of the network into a multi-dimensional feature space Tang *et al.* (2015); Perozzi *et al.* (2014); Grover and Leskovec (2016); Hamilton *et al.* (2019) so that the link prediction problem can then be readily solved in the feature space by using the vast of-the-shelf machine learning algorithms. In the context of social network analysis, there exists a plethora of papers and effective methods for network embedding or graph representation learning Perozzi *et al.* (2014). Most of these embedding techniques rely on a basic premise that nodes that are “similar” in the graph are also “close” in the embedding (latent) space. Currently, the dominant algorithms for the node embedding utilize the random walk-based objectives to define the node level “similarity" Tang *et al.* (2015); Perozzi *et al.* (2014); Grover and Leskovec (2016); Hamilton *et al.* (2019) and train an encoder to satisfy this objective, or even this objective is further simplified to the reconstruct of adjacency matrix via embedding vectors’ dot product decoder, known as Graph Auto-Encoder (GAE) Kipf and Welling (2016b); Gilmer *et al.* (2017).

While powerful, random walk methods suffer from some known limitations, such as degree biasness Erten *et al.* (2011); Coskun and Koyutürk (2015); Stanfield *et al.* (2017), the over-emphasizing of proximity information at the expense of structural information Ribeiro *et al.* (2017), and hyperparameter choice dependent performance Perozzi *et al.* (2014); Grover and Leskovec (2016). To circumvent the random walk-based objective’s limitations in the node embedding context, very recently, Veličković *et al.* (2019) propose a method, Deep Graph Infomax (DGI), that aim at learning a node embedding that maximizes the mutual information between embedding vectors and the global representation of the entire graph. To this end, DGI Veličković *et al.* (2019) trains a single-layered *Graph Convolutional Network* (GCN) encoder Kipf and Welling (2016a). Owing the mutual information maximization objective, DGI has three appealing properties: first, it naturally integrates node features into network embedding by utilizing the single-layered GCN encoder Kipf and Welling (2016a), second, it is trained fully unsupervised manner without requiring node labels, and third, it captures global structure of the network by learning embedding vectors global behaviours. Despite DGI’s Veličković *et al.* (2019) effectiveness, it still uses the single-layered GCN encoder Kipf and Welling (2016a) which completely relies on a random walk-dynamic via symmetrically normalized Laplacian matrix (a.k.a, Laplacian convolution matrix).

In this paper, for the first time in the GCN literature, within the DGI Veličković *et al.* (2019), we propose to integrate various node similarity measures into the single-layered GCN encoder as convolution matrices and learn the network embedding along this line. To this end, we explore the effectiveness of the node similarity measures as convolution matrices in the single-layered GCN encoder by selecting 8 representative local measures Zhou *et al.* (2009) and obtain their graph embedding via DGI. Taking three important *biomedical link prediction tasks*: DDA prediction Gottlieb *et al.* (2011), DDI prediction Zhang *et al.* (2018), and PPI prediction Wang *et al.* (2017) as benchmark problems, we show that, in DGI Veličković *et al.* (2019), these local proximity measure-based GCN encoders (i.e, for all 8 local measures) always outperform the traditional Laplacian convolution matrix-based GCN encoder when the underlying graph has a density that is above a certain threshold and its Laplacian matrix’s eigenvalue decreases sharply which is the case for most of the real-world networks Veličković *et al.* (2019); Kipf and Welling (2016a). Interestingly, we observe that the traditional single-layered GCN encoder with Laplacian is only useful when the network is very sparse and its eigenvalue decay is slow. Although the proposed framework is simple, it opens a rich space for exploration since it completely eliminates the random-walk dynamics, by eliminating to date usage of Laplacian convolution matrix, which is the first step of Random Walk with Restarts measure, from the GCN encoder and we expect the idea brings many useful proprieties into GCN literature.

## 2 Methods

In this section, we first define biomedical link prediction problem and how it can be solved via graph embedding methods. Next, we present a brief background on two graph convolution network (GCN) Kipf and Welling (2016a) based approaches, namely Graph Autoencoder (GAE) Kipf and Welling (2016b) and Deep Graph Infomax (DGI) Veličković *et al.* (2019) and how they aim to solve network embedding problem by setting different objective functions. We then present an overview about how these two embedding techniques, GAE and DGI, facilitate Laplacian convolution matrix in their GCN encoders Kipf and Welling (2016b); Veličković *et al.* (2019). Subsequently, we show that this Laplacian convolution matrix within the GCN encoder relies on random walk dynamic. To avoid this reliance and to comprehensively characterize the effect of the convolution matrix on algorithm performance for link prediction in biomedical networks, we propose to utilize network similarity measures as convolution matrices.

### 2.1 Link Prediction and Network Embedding

The task of predicting new links that are likely to emerge/disappear in a network is a fundamental problem in network analysis (Lü and Zhou, 2011). In the context of biomedical networks, link prediction is useful in discovering previously unknown associations or interactions, as well as identifying missing or spurious interactions (Yue *et al.*, 2020). Motivated by widespread application of network models in computational biology, significant effort has been devoted to the development of algorithms for link prediction in various types of biomedical networks, including protein-protein interactions (PPIs) (Wang *et al.*, 2017), drug-drug associations (Liang *et al.*, 2017), drug-disease associations (Zhang *et al.*, 2018), disease-gene associations (Erten *et al.*, 2011), and drug response in cancers (Stanfield *et al.*, 2017).

In a general setting, the link prediction problem can be stated as follows: Given a network 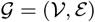, where 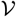 denotes the set of *n* entities (e.g., genes/proteins, biological processes, functions, diseases, drugs etc.) and *ε* denotes a set of *m* interactions/associations among these entities, predict pairs of entities that may also be interacting or associated with each other (Yue *et al.*, 2020). Link prediction can be supervised or unsupervised, where unsupervised link prediction aims to directly score and rank pairs of nodes using features derived from network topology. Supervised link prediction, on the other hand, uses a set of “training" edges and non-edges to learn the parameters of a function that relates these topological features to the likelihood of the existence of an edge.

To extract features that represent network topology, graph representation learning techniques are used to embed the nodes of the network into a multi-dimensional feature space. For a given biomedical network 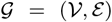, a network embedding is defined as a matrix H ∈ ℝ^*n*×*d*^, where 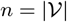 and *d* is a parameter that defines the number of dimensions in the embedding space. Each row of this matrix represents, for each biomedical entity 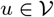, the embedding of *u* as h_*u*_ ∈ ℝ^*d*^.

To facilitate supervised link prediction using node embeddings as features, a given number of edges are randomly sampled from ε. To generate a set of “negative" samples, the same number of node pairs from the set 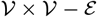 are also randomly sampled. Next, for a given pair of nodes 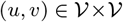, their corresponding embeddings, h*u*, h*v* ∈ ℝ^*d*^ are concatenated with the label 1 or 0 depending on whether (*u, v*) represents a positive (extant edge) or negative (non-extant edge) sample. Finally, these combined latent features with their labels are fed into a supervised machine learning algorithm (e.g. support vector machine (SVM), Random Forest), to train a classifier for link prediction (Yue *et al.*, 2020).

In a recent study, Yue *et al.* (2020) extensively investigate the effectiveness of network embedding techniques in the context of *supervised* link prediction on a broad range of biomedical networks. Their results show that the accuracy of link prediction depends highly on the technique used to compute network embeddings and different embedding techniques can be effective for different applications and networks. Among various embedding techniques, graph convolutional network (GCN) based embedding approaches deliver encouraging results for most of the biomedical link prediction tasks (Yue *et al.*, 2020).

### 2.2 Network Embedding via Graph Convolutional Networks

Graph Convolutional Networks (GCNs) are simplified versions of Graph Convolutional Neural Networks (GCNNs), which are generalizations of conventional Convolutional Neural Networks (CNNs) on graphs (Li *et al.*, 2018). In the context of various machine learning tasks, GCNs are used to facilitate use of network topology in computing latent features from input features associated with network nodes. GCNs are also used to compute node embeddings, i.e., features that represent network topology, by setting the loss function appropriately to capture the correspondence between the embeddings and network topology.

In GCNs, each graph convolution layer involves three steps: 1) feature propagation, 2) linear transformation, and 3) application of a non-linear activation function (Wu *et al.*, 2019). Feature propagation is accomplished by using a convolution matrix that is computed from graph topology. In the context of computing network embeddings, the choice of convolution matrix is critical as it defines the relationship between network topology and computed embeddings. The parameters of linear transformation are learned by training the GCN to minimize loss function and standard non-linear functions are used for activation (.e.g, sigmoid or ReLU). Thus, the key ingredients of a GCN-based network embedding technique are the choice of the convolution matrix and the loss function.

#### Graph Autoencoder (GAE)

Kipf and Welling (2016b) propose GAE as a direct application of their GCN model (Kipf and Welling, 2016a) to the computation of node embeddings in a network. GAE uses the graph Laplacian as the convolution matrix in a two-layer neural network. In this context, the graph Laplacian is defined as

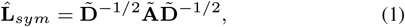

where **A** is the adjacency matrix of the network 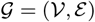, **Ã** = **I** + **A** is the adjacency matrix with self loops added, D = *diag*(*d*_1_, *d*_2_, …, *d*_*n*_) is the degree matrix, and 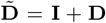. Using 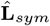 as the convolution matrix, GAE defines the network embedding matrix **H**_*GAE*_ as:

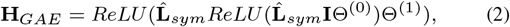

where Θ^(0)^ and Θ^(1)^ are trainable weight matrices. These weight parameters are trained using the following loss function:

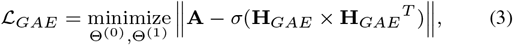

where *σ* denotes logistic sigmoid function.

#### Deep Graph Infomax (DGI)

Veličković *et al.* (2019) develop DGI using the infomax principle (Linsker, 1988) to define a loss function that can be used in various learning settings. In the context of neural networks, DGI computes the embedding matrix **H**_*DGI*_ using a single layered neural network:

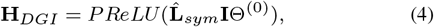

where PReLU denotes parametric ReLU (Veličković *et al.*, 2019) as the non-linear activation function and Θ^(0)^ is a trainable weight matrix. The loss function used to train Θ^(0)^ is defined as binary cross entropy loss:

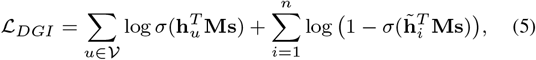

where 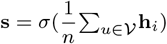 represents the global graph-level summary, 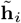 for 1 ≤ *i* ≤ *n* denote the corrupted embedding vectors that are obtained by shuffling the nodes (randomly permuting the rows of **I**), and **M** ∈ ℝ^*d*×*d*^ is a trainable scoring matrix.

Although DGI has not yet been implemented in biological applications, it has demonstrated great potential in other applications Veličković *et al.* (2019). DGI owes its promising results to two factors: (i) capturing the global information of the network by incorporating node summaries and corrupted embeddings in its loss function, and (ii) utilizing the power of this loss function to reduce the number of layers, thereby the number of parameters to be learned. However, the single-layered nature of DGI also limits its ability to diffuse information across the network. In the context of link prediction, node embeddings are utilized to assess the similarity between pairs of nodes. Motivated by this consideration, we hypothesize that coupling of DGI’s neural network architecture and loss function with convolution matrices that are based on node similarities can deliver superior link prediction performance as compared to convolution matrices that directly incorporate the adjacency matrix of the network.

### 2.3 Node Similarity Measures as Convolution Matrices

Algorithms for GCN-based network embedding demonstrate great promise in link prediction (Kipf and Welling, 2016a; Veličković *et al.*, 2019; Hamilton *et al.*, 2019; Wu *et al.*, 2019; Coskun, 2019). Effective application of these methods to link prediction tasks in biology requires careful consideration of the design choices that are encountered in the context of a specific problem. An important design choice in GCN-based network embedding is the choice of the convolution matrix. As discussed above, most of the existing algorithms use the Laplacian as the convolution matrix. To date, the effect of the convolution matrix on algorithm performance has not been comprehensively characterized in the context of link prediction in biomedical networks.

The benefit of a single-layered neural network (as in DGI) over a multi-layered neural network (as in GAE) is that, this reduces the number of parameters in the model and avoids over-smoothing. However, it also limits the convolution to a single step in the network. Based on this observation, we stipulate that network similarity measures that capture local network information can be effective as convolution matrices in conjunction with a single-layered neural network. Such measures include those that have demonstrated success in earlier applications of link prediction, including Common Neighbors, Adamic-Adar, and others (Liben-Nowell and Kleinberg, 2007), Below, we describe these measures and discuss how they can be adopted into the framework of DGI as convolution matrices. In other words, we consider the following formulation for computing node embeddings (where Θ^(0)^ is optimized using the loss function in (5)):

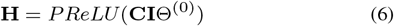

and discuss various options for the convolution matrix C based on the rich literature on unsupervised link prediction. Observe that, for both DGI and GAE, 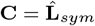.

#### (i) Common Neighbors (CN)

For a given node 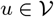, let 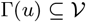 be the set of neighbors of *u*. Then, the count of common neighbors of nodes 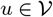 and 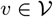 is defined as (Zhou *et al.*, 2009):

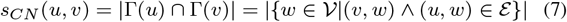

Since (*Â*^2^)_*u,u*_ = *d*_*u*_ + 1 and for *u* ≠ *v*, (*Â*^2^)_*u,u*_ = *s*(*u, v*), the convolution matrix representing count of common neighbors can be computed as:

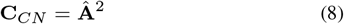

#### (ii) Jaccard Index

This measure assesses the overlap between two sets by normalizing the size of their intersections by the size of their union. In the context of link prediction, Jaccard Index is used to assess the degree of overlap between the neighbors of two nodes in a network Zhou *et al.* (2009):

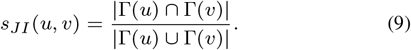

In matrix form, Jaccard Index can be formulated as a convolution matrix as follows:

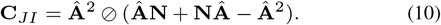

Here, N denotes an all-ones matrix with the same size as A and ⊘ denotes element-wise (Hadamard) division.

#### (iii) Adamic-Adar (AA)

This commonly utilized measure of node similarity refines the notion of common neighbors by assigning more weight to less-connected common neighbors (Adamic and Adar, 2003):

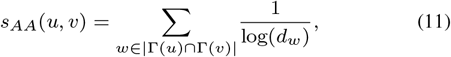

where *d*_*w*_ denotes the degree of node 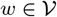. This notion of node similarity can be formulated as a convolution matrix as (Liben-Nowell and Kleinberg, 2007):

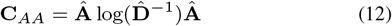

#### (iv) Resource Allocation (RA)

As Adamic-Adar, this measure also aims to reduce the effect of highly-connected common neighbors, but does so more agressively by normalizing with the degree of the neighbor (Zhou *et al.*, 2009). Thus Resource Allocation based convolution matrix can be formulated as:

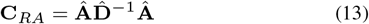

#### (v) Hub Depressed Index (HDI)

Similar to Jaccard Index, HDI aims to normalize the overlap between neighbors of two nodes based on the degrees of the nodes, but does so by focusing on the node with higher degree Zhou *et al.* (2009), i.e.:

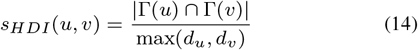

Using the notation introduced above, HDI-based convolution matrix can be formulated as:

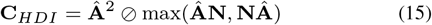

#### (vi) Hub Promoted Index (HPI)

In contrast to Hub-Depressed Index, HPI normalizes the size of the overlap of the neighbors of two nodes by the degree of the less-connected node, thereby promoting hubs. This index can be represented as a convolution matrix as:

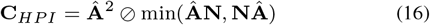

#### (vii) Sørenson Index (SI)

Similar to Jaccard Index, SI normalizes the size of the overlap of the two nodes by taking into account the degree of the two nodes, but uses the sum of the degrees instead of the size of the union:

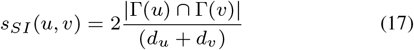

Thus, compared to JC, SI is more conservative toward high-degree nodes as the common neighborhood is counted twice in the denominator. SI can be formulated as a convolution matrix as follows:

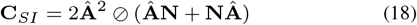

#### (viii) Salton Index (ST)

ST also normalizes the size of the overlap by the degrees of the two nodes, but uses the geometric mean instead of the arithmetic mean for this purpose:

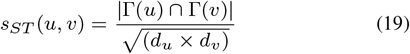

ST can be formulated as a convolution matrix as follows:

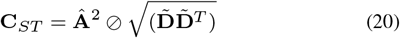

where the square-root operation is applied element-wise to the *n* × *n* matrix 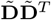.

To summarize our approach, given the adjacency matrix Â of a network, we use the node similarity measures to compute node embeddings for all nodes in the network as follows:

1. Compute the convolution matrix C based on the specified node similarity index (CN, JI, AA, RA, HDI, HPI, SI, or ST).
2. Compute embeddings **H** = *PReLU*(CIΘ^(0)^) using the single-layered GCN encoder
3. Randomly row-wise shuffle I to obtain corrupted node identities Î.
4. Compute corrupted embeddings 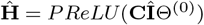, using the single-layered GCN encoder.
5. Compute network-level summary of embeddings 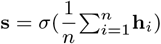
6. Update Θ^(0)^ and M using gradient descent to minimize 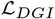 (5).

Once the node embeddings are computed using the above (unsupervised) procedure, we feed these embeddings into BIONEV, the supervised link prediction algorithm implemented by Yue *et al.* (2020).

## 3 Results

In this section, we systematically compare the performance of graph Laplacian vs. node similarity measures as convolution matrices for graph embedding in the context of various link prediction tasks in biology. We start our discussion by describing the datasets and the experimental setup. We then compare the performance of Laplacian and node similarity measures using Area Under Curve (AUC) as a performance criterion for link prediction. Subsequently, we investigate the underlying reasons for the differences in the performance of these methods. In particular, we investigate the effect of graph density (which can be interpreted as the size of training data) and eigenvalue decay on the performance of the methods. Finally, we compare the link prediction performance of the best performing node similarity based convolution matrices against nine other state-of-the-art embedding methods.

### 3.1 Datasets and Experimental Setup

In our experiments, we use four biomedical networks compiled by Yue *et al.* (2020). The statistics of these four networks are shown on Table 1. These networks represent link prediction tasks in the context of three different biomedical applications:

- **DrugBank Drug-Drug Interactions (DDIs)**: The DrugBank-DDI network is composed of verified pairwise interactions between chemical compounds used as drugs, obtained from DrugBank (Wishart *et al.*, 2018). DrugBank is a freely accessible online database that contains detailed information about drugs and drug interactions (Yue *et al.*, 2020).
- **CTDDrug-Disease Associations (DDA)**: Comparative Toxicogenomics Database (CTD) is a database that catalogues the effects of environmental exposures. It contains associations between chemicals and diseases, representing toxic effects of chemicals (Davis *et al.*, 2019). We refer to this dataset as the CTD-DDA network.
- **NDFRT Drug-Disease Associations (DDA)**: This dataset contains drug-disease associations based on National Drug File Reference Terminology (NDFRT) in the unified medical language system, in which there is an edge between a disease and drug if the drug is used for the treatment of the disease (Bodenreider, 2004). We refer to this dataset as the NDFRT-DDA network.
- **Protein-Protein Interactions (PPIs)**: The PPI network contains *Homo sapiens* PPIs extracted from the STRING database (Szklarczyk *et al.*, 2015).

**Table 1.**
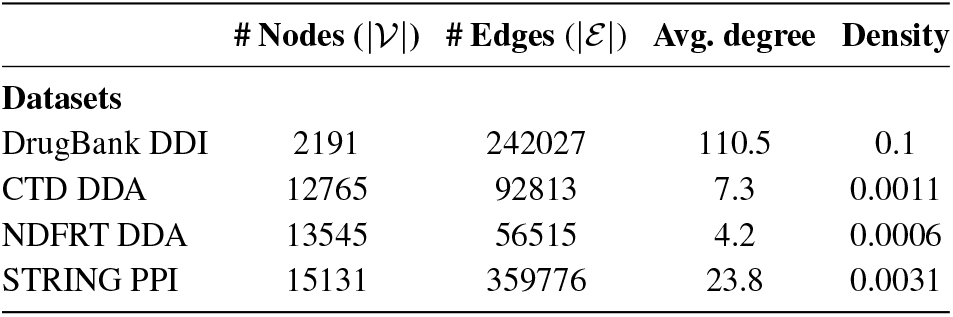
Descriptive statics of the networks used in computational experiments. Avg. degree is defined as 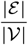, density is defined as 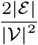

For GAE and DGI algorithms, we use the Python implementation provided respectively by Kipf and Welling (2016a) and Veličković *et al.* (2019). For other state-of-the-art network embedding methods (Table 2), we use OpenNE (https://github.com/thunlp/OpenNE), Python source code implementation. We implement our node similarity measure-based embedding methods on top of PyTorch implementation provided by Veličković *et al.* (2019).

**Table 2.** Caption

| Datasets | Local – NN |  |  | Global – NN |  |  | RandomWalk |  |  | MatrixFactorization |  |  |
| --- | --- | --- | --- | --- | --- | --- | --- | --- | --- | --- | --- | --- |
|  | RA | HDI | HPI | GAE | DGI | LINE | DeepWalk | node2vec | struc2vec | SVD | HOPE | GraRep |
| DrugBank_DDI | 0.912 | **0.924** | 0.918 | 0.891 | 0.896 | 0.909 | 0.917 | 0.889 | 0.901 | 0.912 | 0.910 | 0.913 |
| CTD_DDA | 0.961 | 0.960 | **0.962** | 0.866 | 0.889 | 0.958 | 0.926 | 0.903 | 0.959 | 0.934 | 0.950 | 0.959 |
| NDFRT_DDA | 0.944 | 0.933 | 0.907 | 0.715 | **0.953** | 0.944 | 0.783 | 0.736 | 0.951 | 0.774 | 0.949 | **0.953** |
| STRING_PPI | 0.909 | **0.915** | 0.906 | 0.845 | 0.863 | 0.855 | 0.885 | 0.798 | 0.902 | 0.867 | 0.838 | 0.883 |

We assess the performance of the algorithms using randomized test and training tests, where the randomized tests are repeated 10 times for each algorithm/parameter setting. For each randomized test, we select a certain fraction (referred to as *test ratio*) of the edges in the network uniformly at random, remove these edges from the network, and reserve them as the positive test set. We then compute node embeddings and perform training on the remaining network. For training, we concatenate the embeddings to construct a feature set for each pair of nodes and associated label (1 if the pair has an edge in the training network, 0 otherwise), and use these data to train a Logistic Regression based binary classifier. Subsequently, we make predictions for the pairs of nodes in the positive and negative test sets and compute the area under the ROC curve (AUC) accordingly. For the test ratio, we use %10, %30, and %50 as the fraction of edges removed from the network. For the neural networks used in computing node embeddings, we use default hyper-parameters suggested by the baseline papers and embedding dimension *d* = 100, training epoch 200.

### 3.2 Link Prediction Performance

We compare the link prediction performance of node similarity-based convolution matrices (using a single-layered GCN encoder) against encoders that use the Laplacian matrix for convolution. As representative methods, we use deep graph infomax (DGI, which uses the single-layered GCN encoder we use for the convolution matrices) and graph autoencoder (GAE, which uses a double-layered GCN encoder) for comparison. Selection of these two methods for comparison enables assessment of the effect of the convolution matrix (similarity-based vs. DGI) and the architecture of the GCN (DGI vs. GAE).

Figure 1–4 depict the link prediction performance gain attained when we integrate node similarity measures as convolution matrices into DGI across all datasets for different test ratios, sub-figures a-c. As seen in the figures, node similarity-based methods significantly improve the link prediction performance for all datasets, except NDFRT dataset. These results demonstrate that the node similarity measure-based graph embedding methods are more effective and could be used on various biological link prediction tasks to improve the prediction performance.

**Fig. 1:**
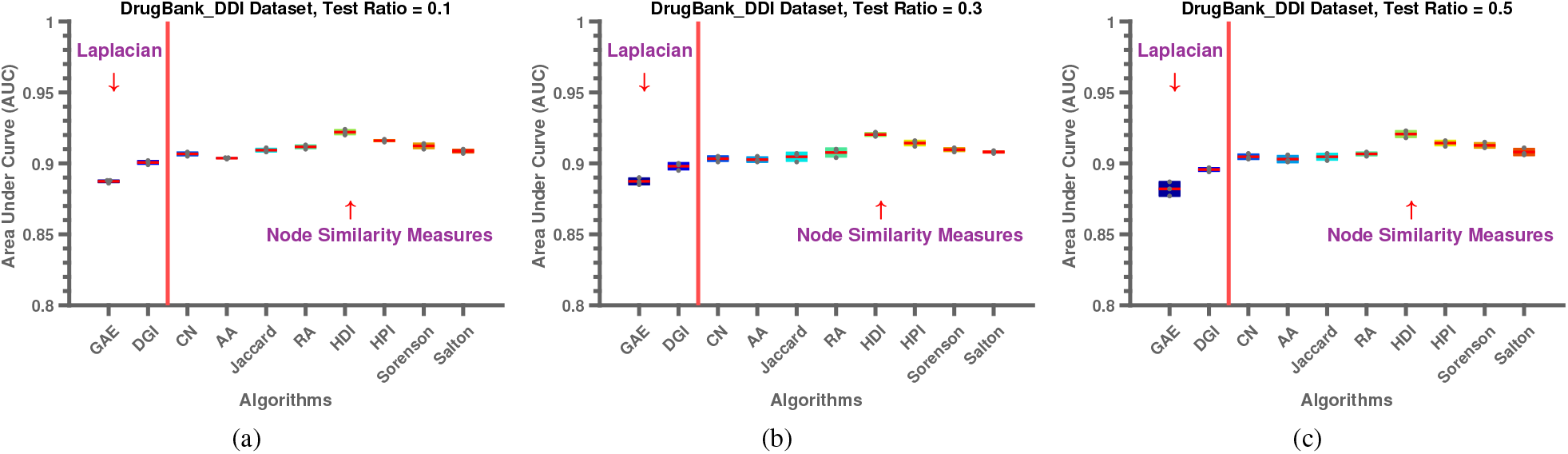
Performance evaluation of. The plots show mean and standard deviation.

**Fig. 2:**
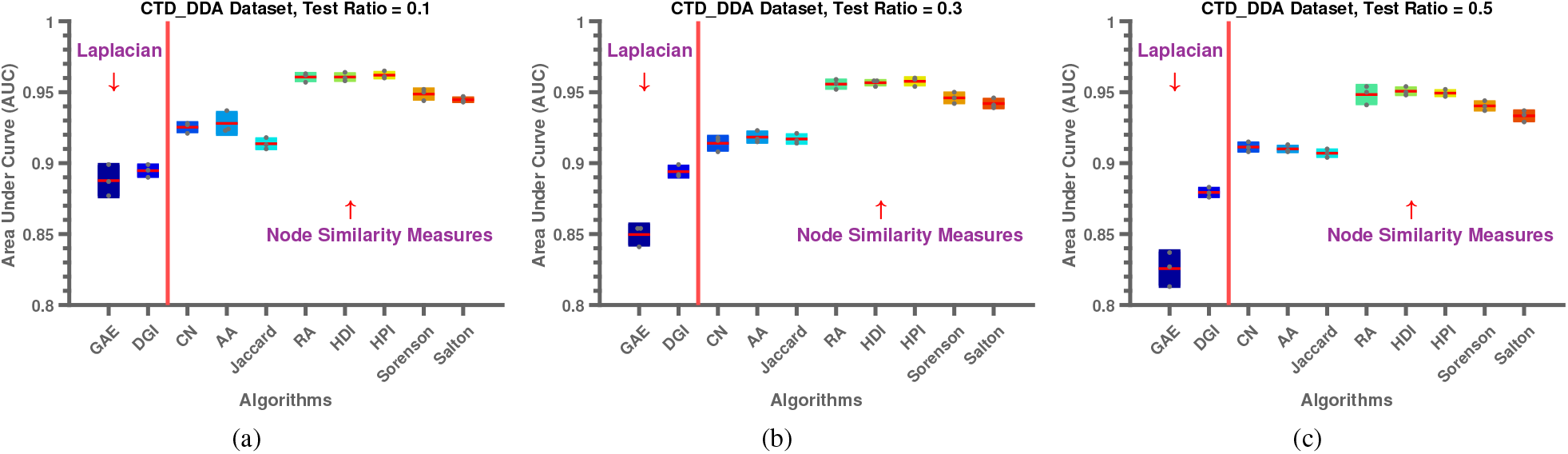
Performance evaluation of. The plots show mean and standard deviation.

**Fig. 3:**
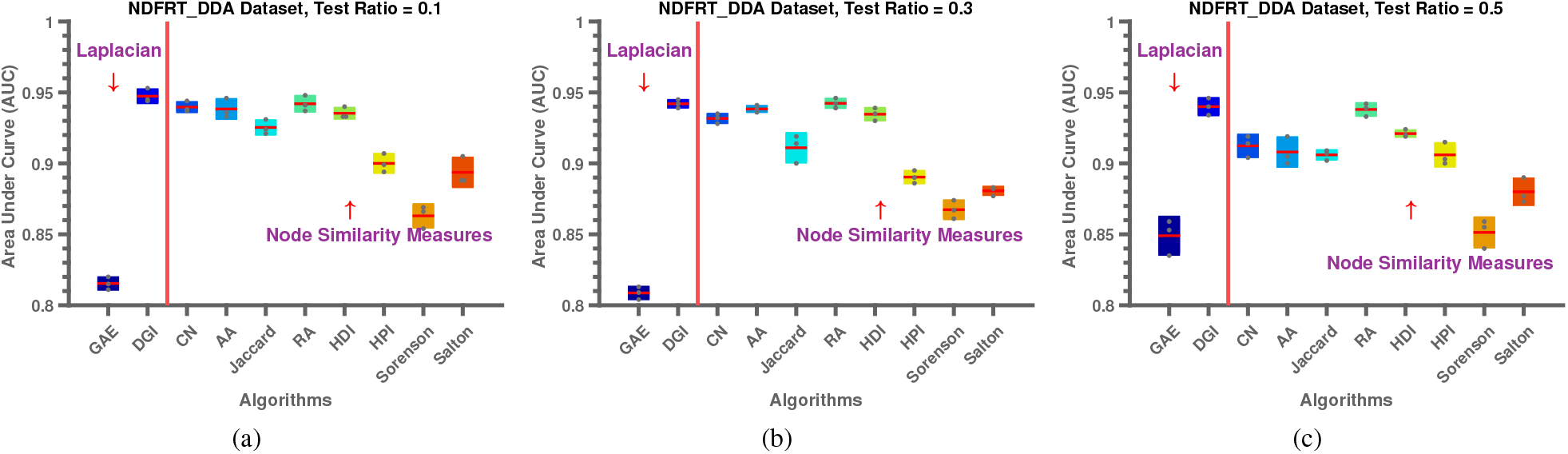
Performance evaluation of. The plots show mean and standard deviation.

**Fig. 4:**
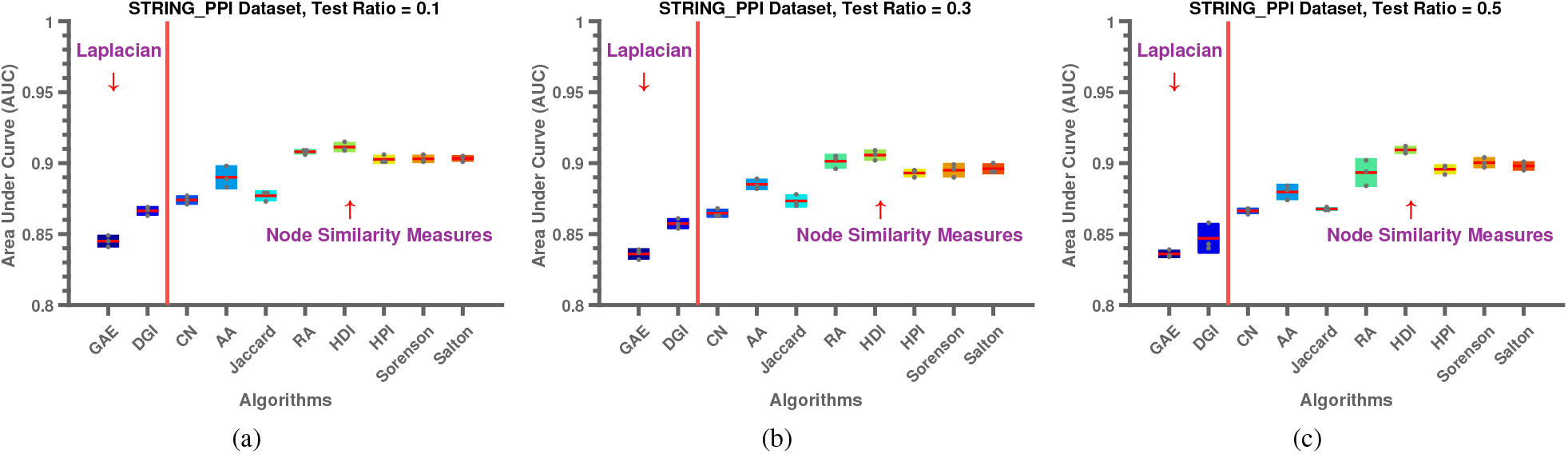
Performance evaluation of. The plots show mean and standard deviation.

### 3.3 When Global Methods Perform Better

Node similarity measures-based algorithms perform substantially better than the baselines on three out of four biomedical networks on link prediction tasks. They perform slightly worse than the baseline on the NDFRT network, which has the least density. To understand this phenomenon, we perform another set of experiments on DrugBank_DDI dataset by gradually decreasing its density. More specifically, in this experiment, we remove the half of existing edges on DrugBank_DDI to reduce its density from the original density, 0.1, to 0.0005. We then perform node embedding with the two best performing node similarity-based single-layered GCN encoder, HDI and HPI, and compare their link prediction performance against the DGI Veličković *et al.* (2019) method, which uses Laplacian convolution matrix, on all sparsified datasets. The results of this analysis for all sparsified and original DrugBank_DDI datasets is shown in the Figure 5(a). As seen in the figure, the network’s density plays a major role on the performance of node similarity-based methods. We further investigate the decay of top eigenvalues of self-loop added Laplacian matrix to shed lights on the phenomenon. To this end, we show the eigenvalue decay of the sparsified networks in Figure 5(b) and all datasets in Figure 5(c). Interestingly, density and top eigenvalue decay exhibit very similar trend suggesting that if the underlying network density is very small and/or eigenvalue decay of its Laplacian matrix is slow, then the Laplacian convolution matrix may be employed for embedding; otherwise node similarity-based GCN encoders are much better alternative to it for the link prediction tasks.

**Fig. 5:**
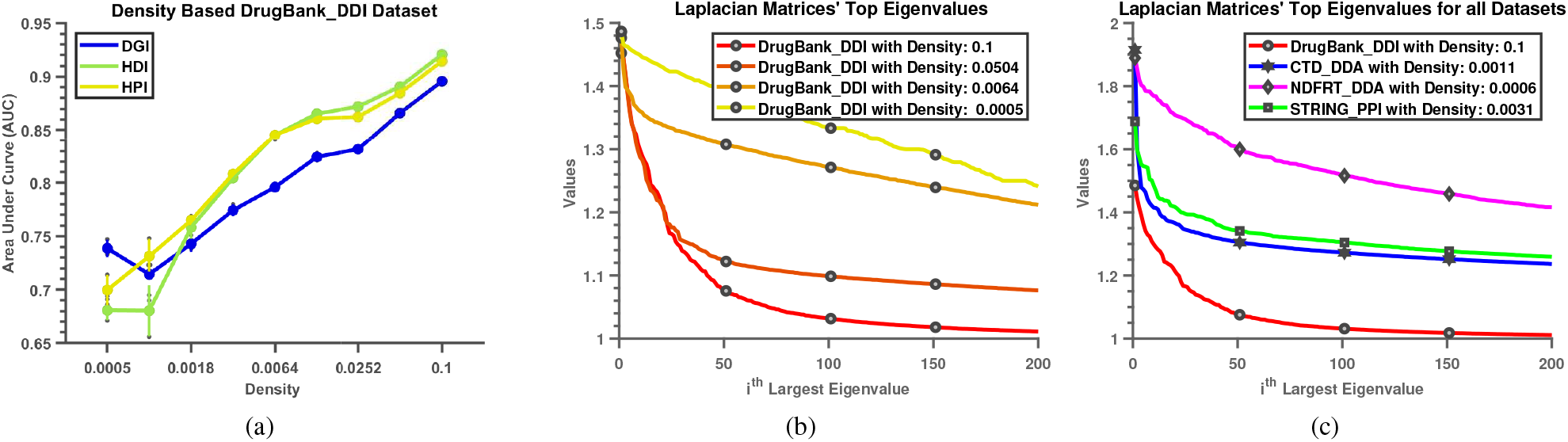
Performance evaluation of. The plots show mean and standard deviation.

### 3.4 Compression with other Baseline Methods

To further evaluate the performance of the node similarity-based methods against state-of-the-art methods on link prediction, we select the best three methods reported in Yue *et al.* (2020) from each three different categories (Neural Network-Based, Random Walk-Based, and Matrix Factorization-Based). We report the results of this analysis in Table 2. We underline the best performing methods across all categories. It can be clearly seen that node similarity-based methods mostly outperform all existing network embedding methods on various link prediction tasks. To better highlight the performance gain, we also give a heat-map plot of the best performing method from each categories in Figure 6.

**Fig. 6:**
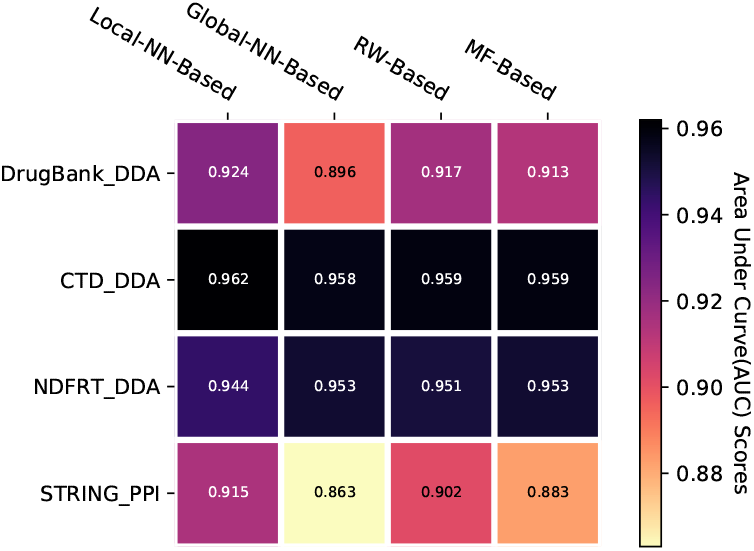
Caption

## 4 Conclusion

In this paper, we propose to use proximity measures as convolution matrices in a single-layered GCN encoder within DGI framework for condensed representation of a given biological network. This new embedding can then readily be used for many down-stream tasks, such as node classification, clustering and link prediction though we, in this paper, particularly focus on link prediction performance of these type of embeddings. Extensive experimental evaluation indicates that node proximity measures in the single-layed GCN encoder deliver much better link prediction results comparing to conventional Laplacian convolution matrix in the encoder. For future, we are planning to learn various proximity measures consensus manner for biomedical and data mining applications.

## Acknowledgements

No text yet

